# Identification of a SARS-CoV-2 host metalloproteinase-dependent entry pathway differentially used by SARS-CoV-2 and variants of concern Alpha, Delta, and Omicron

**DOI:** 10.1101/2022.02.19.481107

**Authors:** Mehdi Benlarbi, Geneviève Laroche, Corby Fink, Kathy Fu, Rory P. Mulloy, Alexandra Phan, Ardeshir Ariana, Corina M. Stewart, Jérémie Prévost, Guillaume Beaudoin-Bussières, Redaet Daniel, Yuxia Bo, Julien Yockell-Lelièvre, William L. Stanford, Patrick M. Giguère, Samira Mubareka, Andrés Finzi, Gregory A. Dekaban, Jimmy D. Dikeakos, Marceline Côté

**Author notes:** To whom correspondence should be addressed: (MC). Contributed equally to this work (Mehdi Benlarbi, Geneviève Laroche).

## Abstract

To infect cells, severe acute respiratory syndrome coronavirus-2 (SARS-CoV-2) binds to angiotensin converting enzyme 2 (ACE2) via its spike glycoprotein (S), delivering its genome upon S-mediated membrane fusion. SARS-CoV-2 uses two distinct entry pathways: 1) a surface, serine protease-dependent or 2) an endosomal, cysteine protease-dependent pathway. In investigating serine protease-independent cell-cell fusion, we found that the matrix metalloproteinases (MMPs), MMP2/9, can activate SARS-CoV-2 S fusion activity, but not that of SARS-CoV-1. Importantly, metalloproteinase activation of SARS-CoV-2 S represents a third entry pathway in cells expressing high MMP levels. This route of entry required cleavage at the S1/S2 junction in viral producer cells and differential processing of variants of concern S dictated its usage. In addition, metalloproteinase inhibitors reduced replicative Alpha infection and abrogated syncytia formation. Finally, we found that the Omicron S exhibit reduced metalloproteinase-dependent fusion and viral entry. Taken together, we identified a MMP2/9-dependent mode of activation of SARS-CoV-2 S. As MMP2/9 are released during inflammation and severe COVID-19, they may play important roles in SARS-CoV-2 S-mediated cytopathic effects, tropism, and disease outcome.

## INTRODUCTION

Coronavirus disease-2019 (COVID-19) is caused by severe acute respiratory syndrome coronavirus-2 (SARS-CoV-2), a highly transmissible positive sense single-stranded RNA virus. The clinical presentation of COVID-19 ranges from asymptomatic or mild to severe disease, including pneumonitis and acute respiratory distress syndrome [1]. Severe COVID-19 is characterized by an uncontrolled release of cytokines, leading to hyperinflammation, tissue damage and dysregulated immune responses. Persistence of these responses often results in multi-organ damage and failure [2]. In addition, upon SARS-CoV-2 infection, pneumocyte syncytia formation is more prevalent in COVID-19 patients with severe chronic respiratory diseases, suggesting a potential hallmark of disease pathogenesis [3].

Membrane fusion is mediated by viral fusion proteins that protrude from the viral membrane or are exposed at the cell surface of infected cells. It can occur during viral entry, or between adjacent cells expressing viral fusion proteins and/or its receptor, causing syncytium formation. To catalyze the merging of membranes during viral entry and cell-to-cell fusion, viral fusion proteins undergo extensive conformational changes, from a high energy metastable state to a highly stable low energy state [4]. This conformational rearrangement is induced by virus specific triggers such as receptor binding, low pH, and/or proteolytic cleavage [4]. For SARS-CoV-2, the viral fusion protein is the spike glycoprotein (S). S is composed of two subunits: S1, which mediates attachment to the host cell receptor angiotensin-converting enzyme 2 (ACE2) and, S2, which facilitates membrane fusion [5]. SARS-CoV-2 and SARS-CoV-1 are related pathogenic betacoronaviruses that share a common host receptor, ACE2, and both require a two-step sequential cellular protease cleavage of the S protein at the S1/S2 junction and at a S2’ site for entry (Fig.1A) [6]. Cleavage of the S1/S2 junction reveals the S2’ site, which is further processed to expose the fusion peptide allowing membrane fusion. However, unlike that of SARS-CoV-1, SARS-CoV-2 S possess an arginine-rich motif within the S1/S2 cleavage site enabling recognition and cleavage at the S1/S2 boundary by furin or furin-like enzymes in the virus-producer cell [7]. The furin cleavage site has been shown to be critical for SARS-CoV-2 infection in human lung cells and transmissibility in ferrets [8, 9]. Moreover, variants of concerns such as Alpha, Delta and more recently Omicron possess mutations within the S1/S2 furin cleavage site that affect furin cleavage efficiency and S fusogenic activity [10–12].

**Figure 1.**
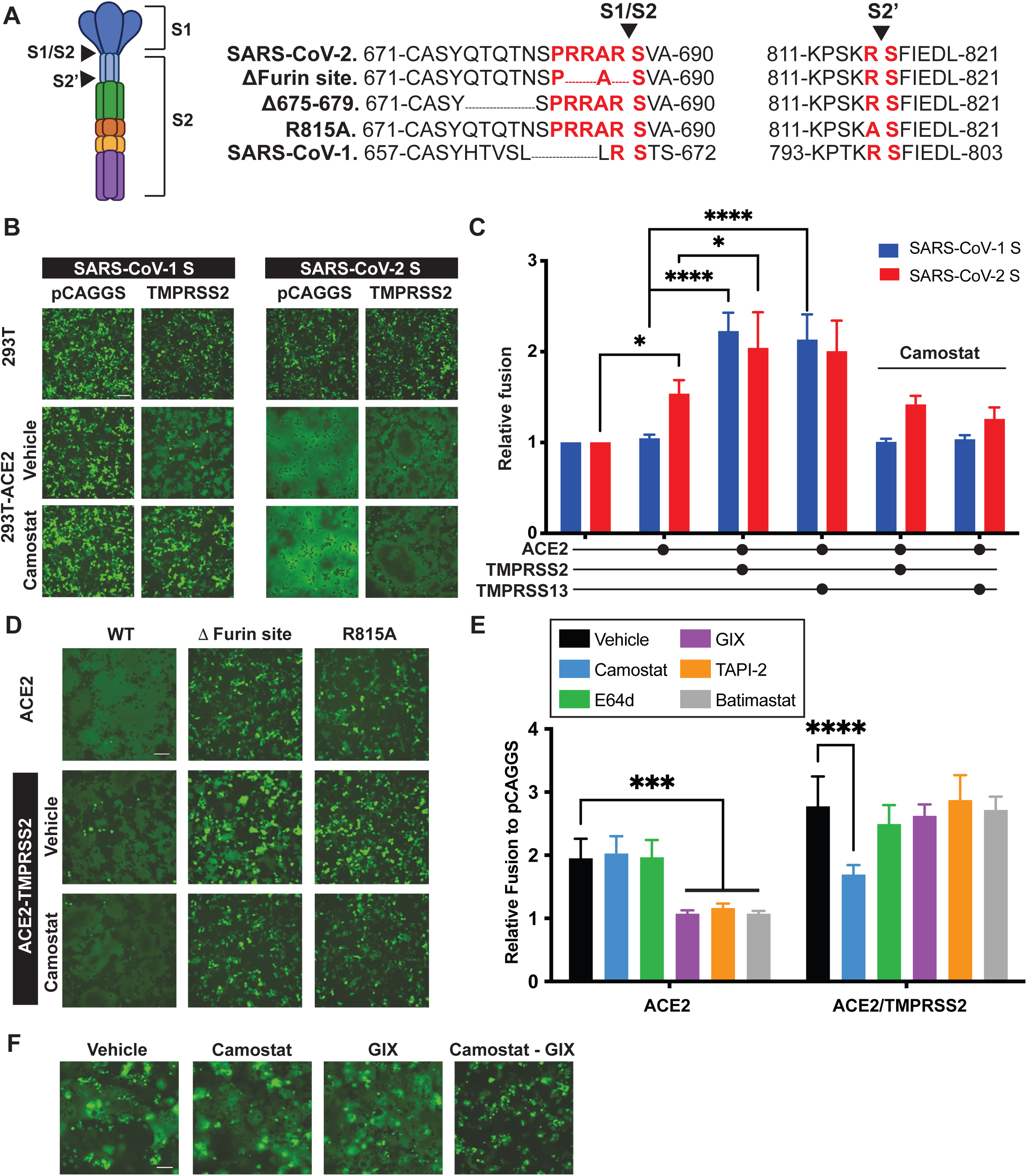
SARS-CoV-2 S can mediate cell-cell fusion in a metalloproteinase-dependent manner. **(A)** Schematics of the S glycoprotein and amino acid sequences at the S1/S2 and S2’ cleavage sites of mutants used in this study. In red, amino acids surrounding the cleavage sites, and arrow heads depict the cleavage site. (Created with BioRender) **(B, D)** 293T or 293T stably expressing ACE2 were co-transfected with plasmids encoding GFP, SARS-CoV-1 or SARS-CoV-2 S WT or indicated mutants, and TMPRSS2, or with an empty vector, in the presence or absence of Camostat (25µM). Syncytia formation was visualized 24 hours post-transfection using fluorescence microscopy. **(C, E)** Effector 293T cells transfected with plasmids encoding SARS-CoV-1 or SARS-CoV-2 S and ZipVenus1, were co-cultured with target 293T cells transfected with plasmid encoding ZipVenus 2, ACE2 and TMPRSS2, TMPRRS13 or empty vector, in the presence or absence of indicated protease inhibitors (Camostat 25µM, E64D 10µM, GIX 25402X3 10µM, TAPI-2 40µM, Batimastat 10µM). Fluorescence generated by the reconstitution of ZIPVenus upon cell-cell fusion was measured at 4 hours of co-culture. **(F)** 293T cells transfected with plasmids encoding GFP and SARS-CoV-2 S were co-cultured (1:1 ratio) with Calu-3 cells in the presence of the indicated inhibitors (Camostat 25µM, GIX 10µM). Syncytia were visualized 24 hours post-transfection using fluorescence microscopy. Each bar shows the mean of triplicate values of 3 independent experiments with error bars showing standard deviation. Significance was determined by analysis of variance (one-way ANOVA) followed by a Dunnett’s multiple comparisons test. P-value lower than 0.05 was used to indicate a statistically significant difference (****, *P* <0.0001, ***, *P* <0.001, **, *P* <0.01, *, *P*<0.05).

Previous studies have defined two possible routes of entry used by SARS-CoV-2 and SARS-CoV-1: an early cell surface pathway following activation by serine proteases, notably the transmembrane serine protease 2 (TMPRSS2), and a late endocytic pathway using endolysosomal cathepsins [13]. Host cell protease expression dictates which viral entry pathways are preferred and could explain why some drugs targeting one but not both pathways are not effective at reducing SARS-CoV-2 burden in patients [14]. S glycoproteins expressed at the surface of infected cells also require similar triggers to induce syncytia formation [15]. Interestingly, previous studies have reported SARS-CoV-2 S-mediated cell-cell fusion in the absence of serine protease expression [16, 17], however the identification of the non-serine protease(s) or if this mechanism represents an additional cell entry pathway remain unknown.

Here we show that in the absence of membrane-bound serine proteases, SARS-CoV-2 S uses the matrix metalloproteinases (MMPs), MMP2/9, to induce cell-cell fusion. In cells expressing high levels of MMP2/9 such as HT1080 cells, infection and syncytia formation induced by replicative Alpha were significantly reduced by MMP inhibitors. We also investigated MMP roles in viral entry using lentiviral pseudotypes and virus-like particles harbouring S of wild-type D614G or variants of concern and found that the various S glycoproteins differentially used the MMP pathway and preferential usage correlated with the extent of S1/S2 processing and syncytia formation. MMPs form a large family of zinc-dependent endopeptidases, most of which, such as MMP2 and MMP9, are secreted [18]. Dysregulation of MMPs have been linked to various human diseases, including cancer, neuronal disorders, and COVID-19 [19–21]. As such, Gelzo, M., et al., reported increased serum levels of MMP3 and MMP9 in severe COVID-19 patients, which also positively correlated with serum interleukin-6 and circulating neutrophils and monocytes [22]. Therefore, in the context of hyperinflammation and dysregulated immune responses, MMPs could play a role in facilitating SARS-CoV-2 viral entry and syncytia formation, expanding tropism to serine protease negative cells, and exacerbating COVID-19. Thus, targeting MMPs, serine proteases, and cathepsins may be useful to reduce SARS-CoV-2 infection and COVID-19 severity.

## RESULTS

### Cell-cell fusion mediated by SARS-CoV-2 S can occur in a serine-protease-independent manner but remains ACE2 dependent and is enhanced by TMPRSS2 and TMPRSS13

To study the host factors required for SARS CoV-2 S activation and compare to host factors required for SARS-CoV-1 S, we first sought to establish a syncytium-formation assay. Parental 293T cells or 293T cells engineered to overexpress human ACE2 were transfected with plasmids encoding S and green fluorescent protein (GFP) to visualize large areas of fused cells that can be distinguishable from single cells. A plasmid encoding for the serine protease, TMPRSS2, or an empty vector, pCAGGS, were also transfected to assess S dependency on serine protease fusion activity. As expected, we found that SARS-CoV-1 S can only induce syncytia formation in the presence of both ACE2 and TMPRSS2 (Fig.1B) [6, 23]. In addition, the ACE2/TMPRSS2-dependent SARS-CoV-1 S cell fusion was abrogated when cells were treated with camostat, a serine protease inhibitor, further indicating that cell fusion by SARS-CoV-1 S requires serine protease activity (Fig.1B). In contrast, SARS CoV-2 S-mediated syncytia were observed even in the absence of TMPRSS2, yet their formation was still dependent on ACE2 expression (Fig.1B). Interestingly, incubation of cells with camostat did not reduce syncytia formation by SARS CoV-2 S, in the presence or absence of TMPRSS2 (Fig.1B). These results suggest that, unlike SARS-CoV-1 S, SARS CoV-2 S-mediated fusion can occur independently of serine protease activity.

ACE2 dependence and the contribution of TMPRSS2 were further assessed by incubating SARS-CoV-2 S-expressing 293T cells with soluble ACE2. In these conditions, an ACE2 dose-dependent increase in S-mediated fusion and a robust enhancement of cell-cell fusion when TMPRSS2 was co-expressed was observed (Fig.1B, S1). To further quantify cell-cell fusion, we used a bimolecular fluorescence complementation assay based on the separate expression of fragments of the yellow fluorescent protein, Venus, fused to a leucine zipper in effector and target cell populations (ZIP Venus assay) (Fig. 1C) [24]. This assay allows for a quantitative measurement of the extent of cell-cell fusion using fluorescence. Previous studies on SARS-CoV-1 S and recent studies on SARS CoV-2 S revealed that other serine proteases, such as TMPRSS4 and TMPRSS13, can also activate these viral fusion proteins [25–29]. Here we sought to validate a role for TMPRSS13 in our assay given its broad expression in the respiratory tract and by immune cells, in addition to its previously reported implications in infection [30]. Target cells transfected with various combinations of plasmids encoding ACE2 and ones encoding TMPRSS2 or TMPRSS13 were co-cultured with effector cells encoding S from SARS-CoV-1 or SARS CoV-2. We found that expression of TMPRSS2 or TMPRSS13 enhanced SARS-CoV-1 and SARS-CoV-2 S mediated cell-cell fusion in an ACE2-dependent manner and that the contribution of TMPRSS2/13 in cell-cell fusion was sensitive to camostat treatment (Fig. 1C). Like the results of the syncytia formation assay, cell-cell fusion was observed in a serine protease-independent manner for SARS-CoV-2 S but not SARS-CoV-1 (Fig. 1C). These results agree with previous studies and indicate that the fusion activity of SARS CoV-1 and SARS-CoV-2 S can be activated by several serine proteases, yet only SARS-CoV-2 can induce cell-cell fusion in a serine protease-independent manner.

### SARS-CoV-2 S2’ site is processed by both serine proteases and metalloproteinases, and cell-cell fusion is abrogated by metalloproteinase inhibitors

Previous studies on coronavirus S glycoproteins showed that activation of the fusion activity of S required sequential proteolytic cleavage at the S1/S2 boundary and at a S2’ site, both of which can be performed by several different serine proteases or endosomal cathepsin proteases during viral entry [31, 32]. To confirm that these protease cleavage steps are also required for the serine protease-independent fusion observed with SARS-CoV-2 S, we next tested the fusion activity of S constructs mutated at the S1/S2 junction (Δ furin site) and the S2’ site (R815A) (Fig. 1A). We found that SARS-CoV-2 S-mediated syncytia formation still occurred when the furin cleavage site was removed, but only in the presence of TMPRSS2 (Fig.1D). In addition, and similar to reports from other studies, a mutation at the S2’ site inactivated the fusion activity of S, which could not be rescued by TMPRSS2 expression (Fig.1D) [17, 33]. These results indicate that the S2’ site of SARS CoV-2 S is critical for fusion activation and that the serine protease-independent fusion requires processing at the S1/S2 boundary.

The requirement for an intact S2’ site also suggested that serine protease independent fusion required proteolysis by an unknown protease. To identify the unknown protease(s) responsible for the fusion, we used the cell-cell fusion assay and incubated the cells with inhibitors of serine proteases (camostat), late endosome/lysosomal cysteine proteases (E64d), and metalloproteinases (GI 254023X (GIX), TAPI-2, and batimastat). As expected, camostat and E64d, did not affect SARS-CoV-2 S-mediated cell-cell fusion in the absence of TMPRSS2. However, all the metalloproteinase inhibitors completely blocked fusion (Fig.1E) suggesting that one or multiple metalloproteinases expressed by 293T-ACE2 cells can trigger SARS-CoV-2 S fusion activity. In addition, these metalloproteinase inhibitors were rendered ineffective when TMPRSS2 was expressed (Fig.1E), suggesting that TMPRSS2 could compensate for an inhibition of metalloproteinase activity. However, metalloproteinase-dependent fusion was still observed in 293T-TMPRSS2/ACE2 cells treated with camostat (Fig. 1D,E). To explore this further, we co-cultured 293T cells expressing SARS-CoV-2 S and GFP with Calu-3 cells which endogenously express ACE2 and serine proteases such as TMPRSS2 [34]. We found that SARS-CoV-2 S efficiently promoted cell-cell fusion which was insensitive to the action of single inhibitors and could only be blocked when a combination of serine protease and metalloproteinase inhibitors were used (Fig.1F). These results indicate that serine proteases and metalloproteinases perform a redundant cleavage step, presumably at the S2’ site.

### SARS-CoV-2 S can mediate cell entry via three distinct routes, including a metalloproteinase-dependent pathway

Previous studies had also reported SARS-CoV-2 S batimastat-sensitive cell-cell fusion, yet this broad metalloproteinase inhibitor had no effect on viral entry in the cell lines tested [16, 17]. However, since our results show that metalloproteinases can activate S for cell-cell fusion, the expectation is that these proteases should also be able to mediate viral entry. To investigate this, we assayed viral entry in 293T-ACE2 and Calu-3 cells using lentiviral pseudotypes. Like other studies [35], we found that SARS-CoV-2 and SARS-CoV-1 S-mediated entry was strongly inhibited by E64d in 293T-ACE2 cells, and by camostat in Calu-3 cells (Fig.2AB). In addition, a slight reduction in entry of SARS-CoV-2 pseudotypes in 293T-ACE2 cells was observed in the presence of metalloproteinase inhibitors (Fig.2A). However, the dramatic decrease in entry after treatment with E64d indicated that the cathepsin entry pathway is the preferred route in these cells.

**Figure 2.**
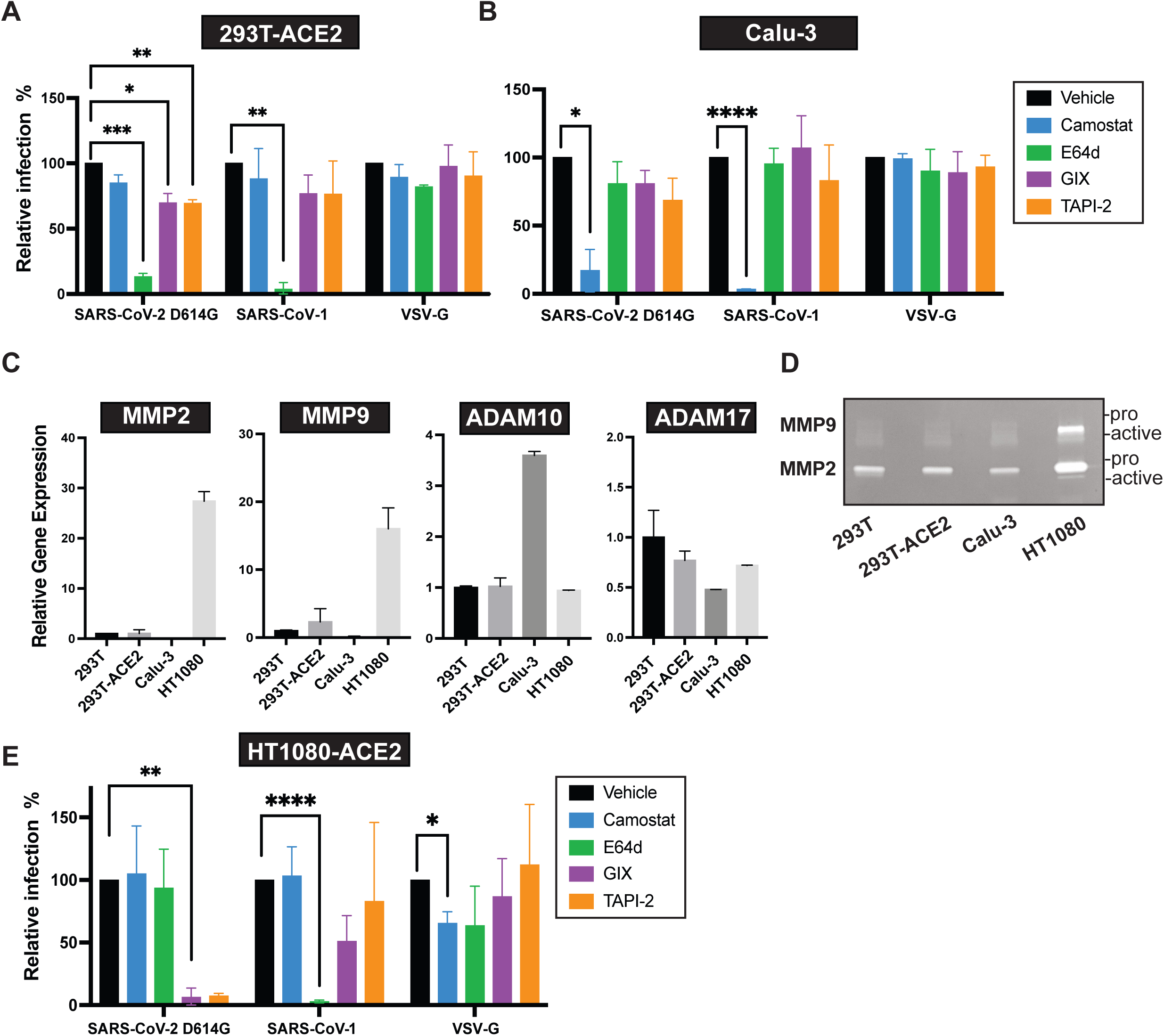
SARS-CoV-2 S can mediate viral entry using three distinct pathways, including a new metalloproteinase-dependent entry route in cells expressing high levels of MMP2 and MMP9. **(A,B,E)** 293T-ACE2, Calu-3 and HT1080 transfected with ACE2, were pre-treated for 1 h with 25 μM Camostat, 10 μM E64D, 40 μM TAPI-2, 10 μM GIX or Vehicle (DMSO) followed by addition of lentiviral pseudoviruses encoding LacZ and bearing the SARS-CoV-2 D614G S, SARS-CoV-1 S, or VSV-G. After 48 h, cells were fixed and stained with X-gal overnight at 37°C and foci representing infected cells were counted. Relative infection was calculated as the number of foci in the indicated inhibitor treatment relative to vehicle treatment. The impact of inhibitors on infection compared to vehicle was analyzed using a two-way ANOVA and Dunnett’s post-hoc analysis. **(C)** Relative mRNA levels of MMP2, MMP9, ADAM10 and ADAM17 in studied cell lines was measured by RT-qPCR. The level of actin mRNA expression in each sample was used to standardize the data, and normalization on 293T gene expression was performed. **(D)** Gelatin zymogram of conditioned media (24h) from indicated cell lines reveals secreted MMP2 (72kDa) and MMP9 (92 KDa) activity, arrows indicate the pro- and active- MMP2 or MMP9. P-value lower than 0.05 was used to indicate a statistically significant difference (****, *P* <0.0001, ***, *P* <0.001, **, *P* <0.01, *, *P*<0.05).

We surmised that while expression of metalloproteinases in 293T-ACE2 cells might be sufficient to induce cell-cell fusion, the levels could be too low to mediate effectively viral entry. We therefore measured metalloproteinase expression in 293T cells and various cell lines using quantitative reverse transcription PCR (RT-qPCR) (Fig.2C, data not shown). We found that HT1080 cells express high levels of MMP2 and MMP9, which are targets of GIX, TAPI-2 (Fig.2C) [36, 37]. The secretion and elevated activity of MMP2 and MMP9 produced by these cells were also confirmed by zymography (Fig.2D). To test a potential role of these proteases in SARS-CoV-2 entry, HT1080 cells were transfected with a plasmid encoding ACE2 and infected with SARS-CoV-2 lentiviral pseudotypes. Strikingly, in these cells, SARS-CoV-2 S-mediated entry was insensitive to E64d and camostat, but completely abrogated by TAPI-2 and GIX. In comparison, entry of pseudotypes bearing SARS-CoV-1 S was blocked by E64d and those with VSV-G remained mostly unaffected, although a slight inhibition by camostat was noted (Fig. 2E). These results indicate that in cells expressing high levels of secreted MMPs, SARS-CoV-2 S entry was mediated via a third, previously unrecognized entry route that is cysteine/serine protease-independent and metalloproteinase-dependent.

### A viral-like-particle system reveals distinct S processing efficiencies and differential usage of the metalloproteinase-dependent entry pathway among SARS-CoV-2 variants

Although lentiviral pseudotypes are suitable, well-established surrogate systems to investigate viral entry, differential budding sites between lentiviruses and coronaviruses can lead to variations of S glycoprotein processing [38, 39]. Since our cell-cell fusion assays clearly demonstrated that processing of S at the S1/S2 junction is required for metalloproteinase-dependent activation of S, we sought to validate the lentiviral pseudotype system using virus-like particles (VLPs). As described previously, SARS-CoV-2 VLPs are produced in cells after co-expression of the four SARS-CoV-2 structural proteins (S, M, E, N) and a reporter gene, thus more accurately recapitulating SARS-CoV-2 egress and entry compared to lentiviral pseudotypes [40]. We found that lentiviruses incorporate fully processed S (Fig.3A), while S on the VLPs were a mixture of unprocessed (S0) and cleaved S (Fig.3B), which is in agreement with previous studies [40–44]. Interestingly, highly transmissible variants of concern Alpha and Delta both have mutations at P681 near the S1/S2 furin cleavage site, however our results demonstrate differential S processing between these two variants [10, 45]. Specifically, the Delta variant S, which harbours a P681R mutation, is processed more efficiently than both the alpha variant S, which harbours a P681H mutation, and the D614G S (Fig.3B). This difference in processing efficiency was not discernible in the lentiviral pseudotype system (Fig.3A). In addition, we tested a previously described deletion mutant (del675-679), often generated during cell culture adaptation of SARS-CoV-2, abrogating the efficiency of S1/S2 processing while keeping the furin cleavage site intact (Fig. 1A) [46–48]. Surprisingly, even this mutant had enhanced processing when expressed on lentiviral pseudotypes compared to VLPs, further demonstrating differential S processing in both systems (Fig. 3A,B).

**Figure 3.**
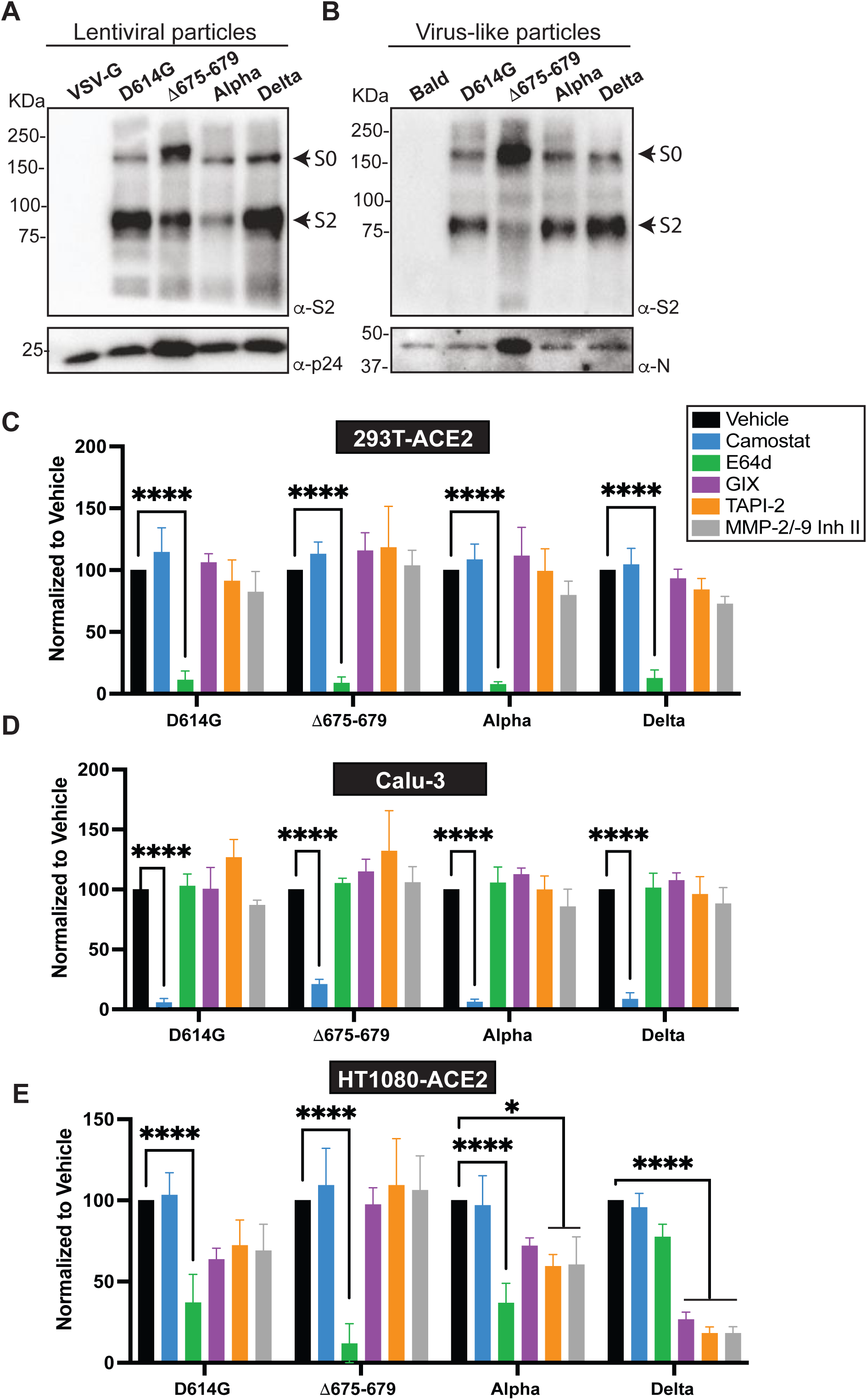
Delta viral-like particles preferentially use the metalloproteinase-dependent entry pathway in HT1080-ACE2 cells. **(A,B)** Processing of spike protein of purified lentiviral (LVP) and virus-like particles (VLP) was analyzed by immunoblot using an anti-S2 antibody allowing the detection of S0 and S2. As for controls, anti-p24 and anti-N antibodies were used for LVP and VLP respectively. **(C,D,E)** VLP entry assay on 293T-ACE2, Calu-3 and HT1080-ACE2 cell pre-treated for 1 h with 25 μM Camostat, 10 μM E64D, 40 μM TAPI-2, 10 μM GIX, 20 μM MMP2/9 inhibitor or Vehicle (DMSO). VLP entry was measured 24h post-infection by measuring the activity of the luciferase reporter. Each bar shows the mean of triplicate values of 3 independent experiments (n=3) with standard deviation. Significance was determined by analysis of variance (one-way ANOVA) followed by a Dunnett’s multiple comparisons test. P-value lower than 0.05 was used to indicate a statistically significant difference (****, *P* <0.0001, ***, *P* <0.001, **, *P* <0.01, *, *P*<0.05).

We next tested entry of SARS-CoV-2 VLPs in 293T-ACE2, Calu-3, and HT1080 cells stably expressing ACE2 (HT1080-ACE2). Similar to the lentiviral pseudotypes, we found that entry of VLPs harbouring S with the D614G mutation or del675-679, or those of Alpha and Delta was strongly inhibited by E64d in 293T-ACE2, and by camostat in Calu-3 cells (Fig.3C,D). However, unlike lentiviral particles, D614G and Alpha VLP entry into HT1080-ACE2 were partially sensitive to E64d, GIX and TAPI-2, while Delta entry was insensitive to E64d and dramatically decreased by GIX and TAPI-2 (Fig.3E). This suggests that Delta S preferentially used the metalloproteinase-dependent pathway. In comparison, del675-679 S-mediated VLP entry was only sensitive to E64d. Given the broad specificity of GIX and TAPI-2, we also tested a specific MMP2/MMP9 inhibitor, MMP-2/-9 Inhibitor II, which phenocopied the broad-spectrum metalloproteinase inhibitors (Fig.3C,D) suggesting an important roles for MMP2 and MMP9 in this pathway. Taken together, these results confirm SARS-CoV-2 entry via a third metalloproteinase-dependent route that is enabled by S1/S2 processing.

### MMP2 and MMP9 knockdown reduces serine protease-independent syncytia formation

The SARS-CoV-2 preferential metalloproteinase entry pathway in HT1080 cells, which express high levels of secreted MMP2/9, in conjunction with the antiviral activity of the MMP-2/MMP-9 inhibitor II, strongly suggests that MMP2 and/or MMP9 play a role in the metalloproteinase-dependent activation of S. To test this, we sought to knockdown expression of MMP2 and MMP9 in 293T-ACE2 and measure syncytia formation. We first validated the knockdown efficiency of three dicer substrate interfering RNAs (dsiRNAs) for both MMP2 and MMP9 in HT1080-ACE2 cells by qPCR and gelatin zymography (Table S1, Fig.S2AB). As expected, production and secretion of both MMP2/9 was efficiently reduced when transfected with their respective dsiRNAs, although to different extents. We chose the two best dsiRNAs for MMP2 and MMP9 respectively for the syncytia formation assay. We transfected 293T-ACE2 cells with dsiRNAs specific for MMP2, MMP9, or scramble dsiRNA as a control, followed by transfection of a plasmid encoding SARS-CoV-2 S to test serine protease-independent fusion. Knockdown efficiency of MMP2 and MMP9 was assessed using gelatin zymography (Fig.4D), although MMP9 was barely visible. Even with incomplete knockdown, we observed a significant decrease in the extent and kinetics of syncytia formation in the MMP2 and/or MMP9 knockdown cells (Fig.4.ABC). Of note, we did not observe a significative effect for one of the MMP9-specific dsiRNA (#1), although significant reduction was seen when in combination with a MMP2-specific dsiRNA (#2). In contrast, there was no difference in syncytia formation for cells transfected with the scramble dsiRNA (Fig.4.AB). These results strongly suggest that MMP2 and MMP9 are playing critical roles in the metalloproteinase-dependent SARS-CoV-2 S-mediated fusion.

**Figure 4.**
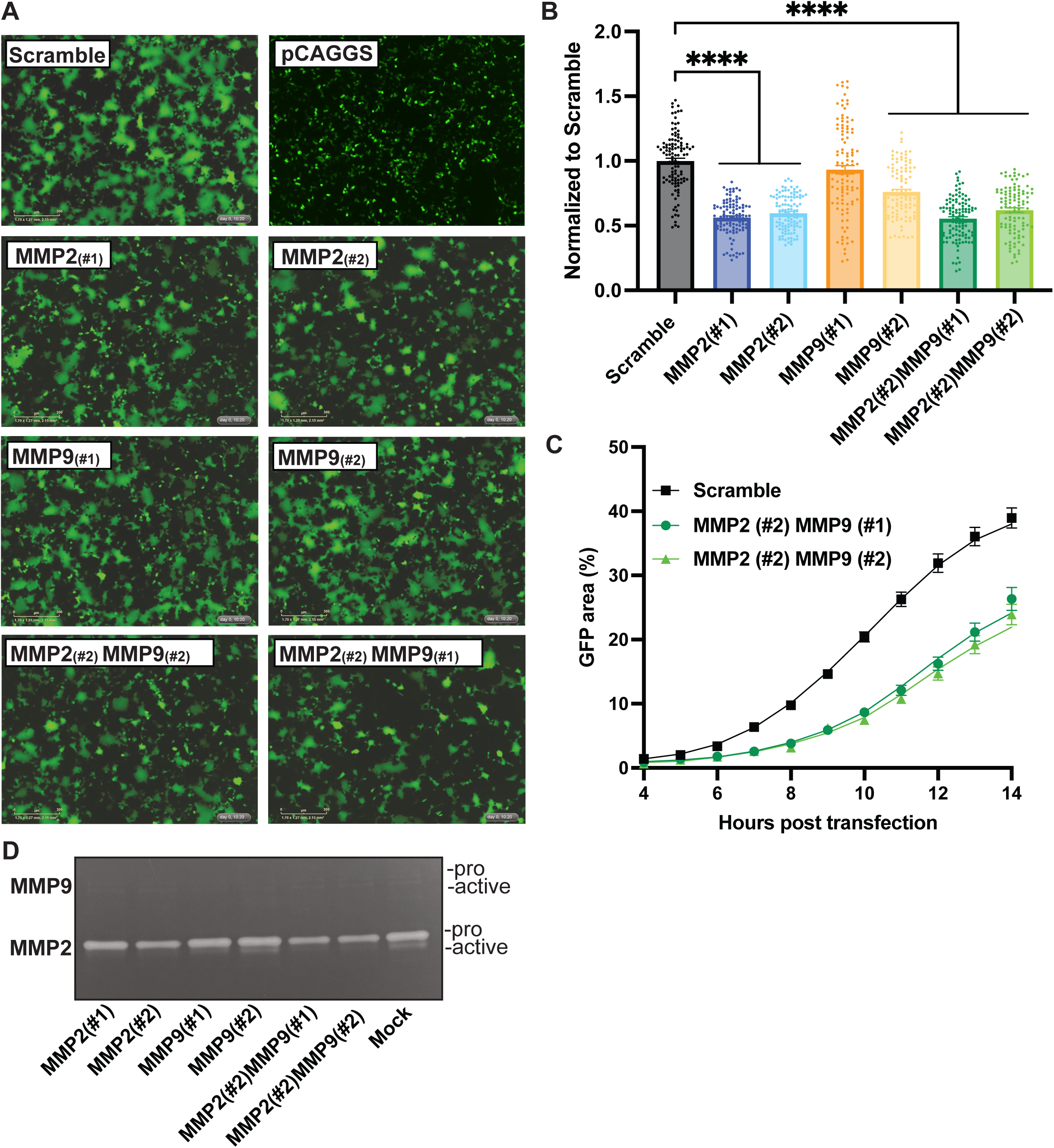
MMP2/MMP9 knockdown reduces the metalloproteinase-dependent syncytia formation. **(A)** Representative images of syncytia formation. 293T-ACE2 cells were transfected with the indicated dsiRNAs alone or in combination (1:1 ratio) at a final concentration of 10nM for 20hours, followed by transfection with pLV-GFP and D614G spike protein. Images were taken 10 hours post-transfection. **(B)** Quantification of GFP+ surface areas at 10 hours post-transfection normalized to scramble dsiRNA. **(C)** Kinetics of syncytia formation. **(D)** Gelatin zymogram of conditioned media (24h) of dsiRNAs transfected 293T-ACE2, arrows indicate the pro- and active- MMP2 or MMP9 Each bar shows the mean of triplicate values of 3 independent experiments (n=3) with standard deviation. Significance was determined by analysis of variance (one-way ANOVA) followed by a Dunnett’s multiple comparisons test. P-value lower than 0.05 was used to indicate a statistically significant difference (****, *P* <0.0001, ***, *P* <0.001, **, *P* <0.01, *, *P*<0.05).

### Metalloproteinase inhibitors block syncytia formation and reduce replication of SARS-CoV-2 Alpha in HT1080-ACE2 cells

We next sought to validate our findings with replicative SARS-CoV-2. Alpha was used to infect HT1080-ACE2 cells overnight in the presence of camostat, E64d, TAPI-2, GIX, or vehicle. The next day, cells were fixed and stained for S protein (S) and nucleocapsid protein (N) to visualize infected cells and an ELISA on N was performed in parallel to quantify infection. We found that Alpha infection of the HT1080-ACE2 cells led to the formation of large multinucleated cells in vehicle, camostat and E64d-treated cells (Fig.5A, S3). While E64d treatment did not prevent syncytia formation, fewer infected cells were observed and N levels were significantly decreased (Fig.5AB, S3). This suggested that cathepsin inhibition decreases infection and viral replication, but not S-mediated cell-cell fusion of infected cells. Similarly, metalloproteinase inhibitors also decreased both the number of infected cells and N expression levels measured by ELISA. However, these drugs also completely blocked syncytia formation (Fig.5AB, S3), indicating that metalloproteinases facilitate both entry and cell-cell fusion. Therefore, in agreement with the VLP assays, in HT1080-ACE2 cells, a proportion of Alpha infections proceeded in a cathepsin-dependent manner, while other infections proceeded in a metalloproteinase-dependent manner. This alternative usage of cathepsin or metalloproteinases is likely linked to the processing efficiency of the incorporated S glycoprotein.

**Figure 5.**
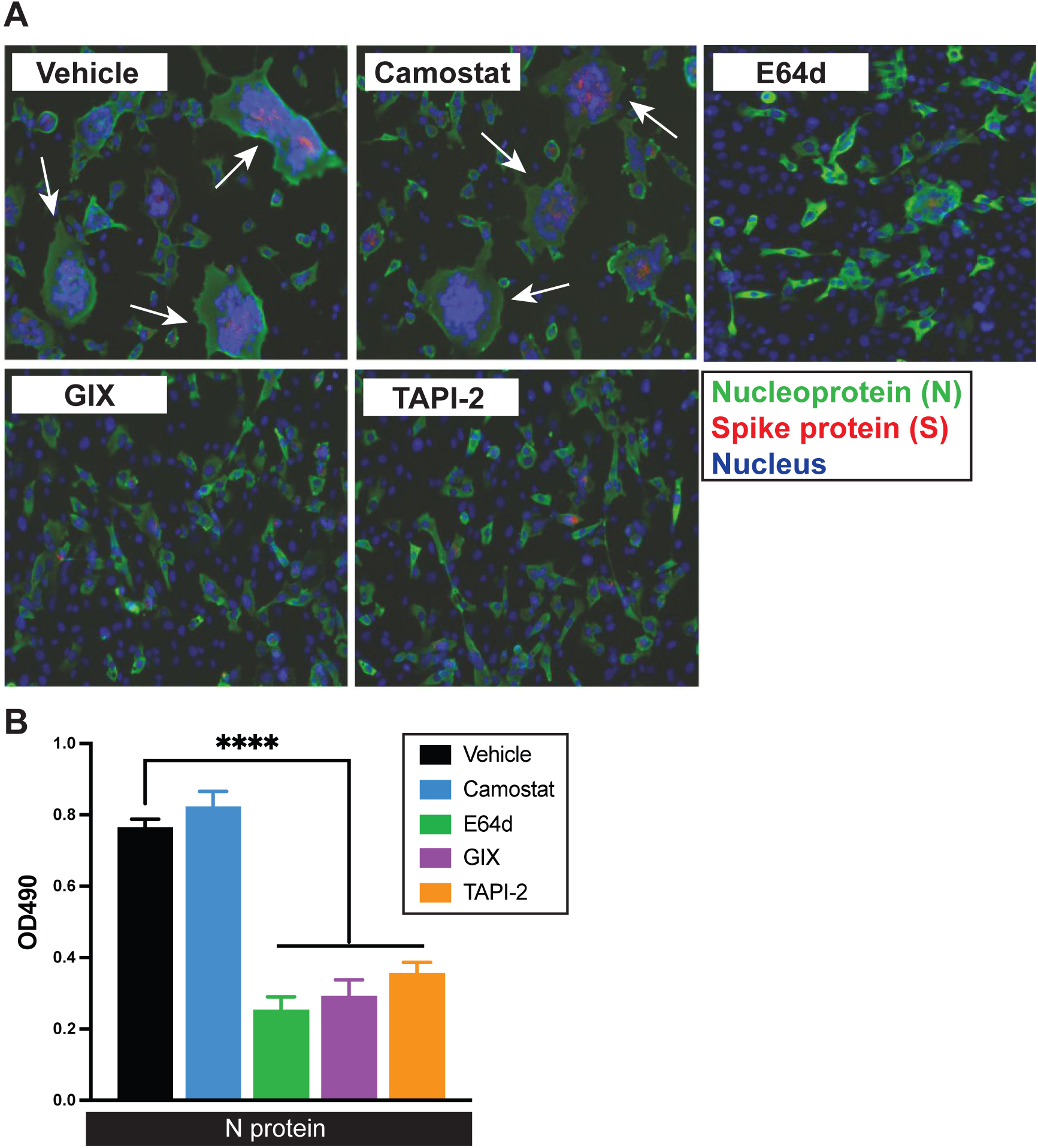
Syncytia formation and Alpha variant infection are blocked by metalloproteinase inhibitors in HT1080-ACE2 cells. **(A**) Visualization of HT1080-ACE2 syncytia formation after infection by Alpha in presence of indicated inhibitors or vehicle. Cells were treated with Camostat (20 uM), E64D (10 uM), TAPI-2 (20 uM), GIX (10 uM) or Vehicle (DMSO) and then infected with Alpha. After 20h, cells were washed, blocked, and stained with rabbit anti-SARS-CoV-2 spike (S), mouse anti-SARS-CoV-2 nucleocapsid (N) followed by staining with DAPI, donkey anti-mouse IgG Alexa Fluor Plus 488 and donkey anti-rabbit IgG Alexa Fluor Plus 594 antibodies. Nuclei, S and N proteins are shown in purple, red and green respectively. Fluorescent images were acquired with an EVOS™ M7000 Imaging System. Images are representative of 3 independent experiments. **(B)** SARS-CoV-2 infection quantification following infection in presence of indicated inhibitors or vehicle. 20h post-infection, cells were washed, blocked, permeabilized and stained with mouse anti-SARS-CoV-2 N protein followed by an anti-mouse IgG HRP in conjunction with SIGMAFAST™ OPD developing solution. Optical density (OD) at 490 nm was measured using Synergy LX multi-mode reader and Gen5 microplate reader and imager software. The red line indicates the OD obtained for vehicle. Each bar shows the mean of triplicate values of 3 independent experiments with error bars showing standard deviation. Significance was determined by analysis of variance (one-way ANOVA) followed by a Dunnett’s multiple comparisons test. P-value lower than 0.05 was used to indicate a statistically significant difference (****, *P* <0.0001, ***, *P* <0.001, **, *P* <0.01, *, *P*<0.05).

### The Omicron S does not efficiently mediate metalloproteinase-dependent cell-cell fusion and preferentially uses the serine protease or cathepsin protease pathways for entry

Since its emergence and discovery, Omicron has spread rapidly worldwide and became the dominant circulating SARS-CoV-2 variant [49]. Interestingly, recent studies have reported decreased processing of the spike glycoprotein and potential reduced activation by TMPRSS2 [11, 50]. Given that our findings support a model by which metalloproteinase usage is dictated by S cleavage at the S1/S2 junction, we sought to investigate whether the Omicron S could be activated in a metalloproteinase-dependent manner. We first performed cell-cell fusion assays using 293T-ACE2 cells as effector cells in the presence or absence of TMPRSS2 expression (Fig.6A). Interestingly, we found that Omicron S mediated only modest fusion in the absence of TMPRSS2, in contrast to the ancestral, D614G, Alpha, or Delta variant S. As expected, Omicron S fusion activity was enhanced in the presence of TMPRSS2, albeit to a slightly lower extent compared to the other SARS-CoV-2 S tested (Fig.6A). In addition, as previously reported and in accordance with our model, the ratio of S2/S0 in cell lysates was reduced for Omicron when compared to D614G (Fig.6B).

**Figure 6.**
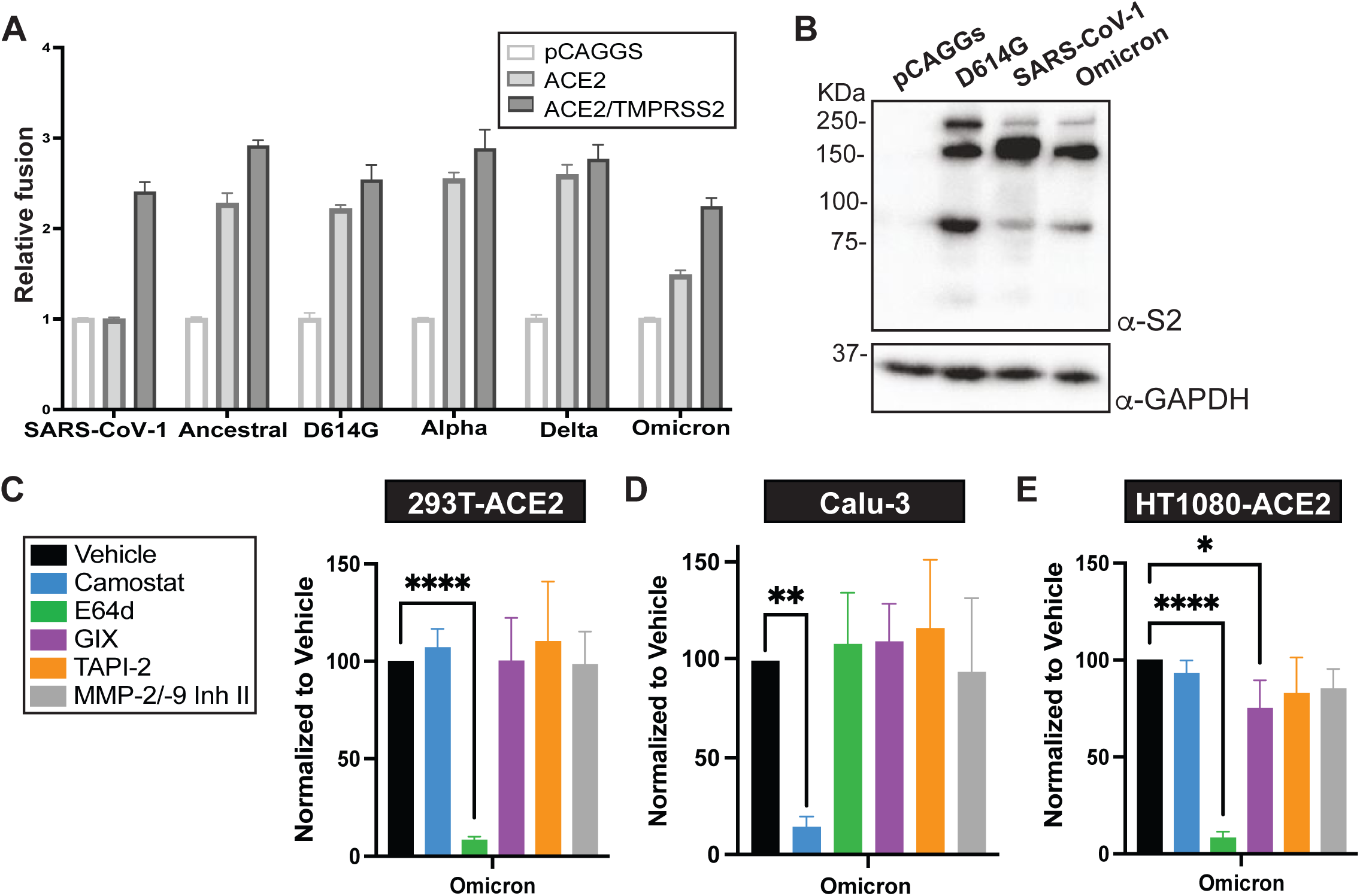
Omicron S is not effectively triggered in a metalloproteinase-dependent manner. **(A)** Effector 293T cells transfected with plasmids encoding the indicated different spikes, and ZipVenus1, were co-cultured with target 293T cells transfected with plasmids encoding ZipVenus 2, ACE2 and TMPRSS2 or empty vector. Fluorescence generated by the reconstitution of ZIPVenus upon cell-cell fusion was measured at 4 hours of co-culture (1:1 ratio). **(B)** Processing of spike protein on effector 293T cells was analyzed by western blot (WB). The S2 protein bands were visualized using an anti-S2 antibody allowing the detection of S0 and S2. As for loading control, anti-GAPDH antibody was used. **(C, D, E)** Omicron Spike VLP entry assay on **(C)** 293T-ACE2, **(D)** Calu-3 and **(E)** HT1080-ACE2 cells pre-treated for 1 h with 25 μM Camostat, 10 μM E64D, 40 μM TAPI-2, 10 μM GIX, 20 μM MMP2/9 Inhibitor or Vehicle (DMSO). VLP entry was measured 24h post-infection by measuring the activity of the luciferase reporter. Each bar shows the mean of triplicate values of 3 independent experiments with error bars showing standard deviation. Significance was determined by analysis of variance (one-way ANOVA) followed by a Dunnett’s multiple comparisons test. P-value lower than 0.05 was used to indicate a statistically significant difference (****, *P* <0.0001, ***, *P* <0.001, **, *P* <0.01, *, *P*<0.05).

We next assessed the preferential entry route employed by Omicron S using VLPs. We found that, similarly to other SARS-CoV-2 variants, Omicron used the endosomal cathepsin-dependent pathway in 293T-ACE2 cells and the surface, serine protease-dependent pathway in Calu-3 cells (Fig.6C). However, strikingly, Omicron did not use the metalloproteinase-dependent pathway in HT1080-ACE2 cells and strictly used the endosomal route (Fig.6C). Further, compared to Alpha, D614G and Delta VLP, Omicron used the TMPRSS2-dependent pathway in Calu-3 cells less efficiently, but had similar entry efficiency to D614G in 293T-ACE2 and HT1080-ACE2 cells (Fig.S4). Taken together, our data suggest a shift in protease usage by Omicron, which may play a part in the differential tropism and pathogenicity observed for this variant.

## DISCUSSION

Previous studies have shown that SARS-CoV-1 and SARS-CoV-2 can enter cells via two distinct ACE2-dependent pathways: a serine protease and a cysteine protease pathway [5, 6]. Here we show that the SARS-CoV-2 wild-type and variant of concerns (VOCs) S glycoprotein can be triggered via a third mechanism that is dependent on secreted metalloproteinases (MMPs), specifically MMP2 and MMP9, for cell-cell fusion and viral entry. The ability to use this entry pathway required high expression of these proteases and proteolytic processing at the S1/S2 boundary in viral producer cells. Accordingly, usage of this pathway by SARS-CoV-2 variants correlated with differential extents of S processing in viral producer cells; the S of Delta preferentially entered via the metalloproteinase route when available, while the S of Omicron did not. Given that metalloproteinases such as MMP2/9 are released and highly expressed in the context of lung damage and inflammation during severe COVID-19, this mechanism of activation could play critical roles in S-mediated cytopathic effects, tropism, and overall pathogenesis.

With previous work reporting the presence of syncytia in the lung of deceased COVID-19 patients [42, 51], the ability of SARS-CoV-2 S to mediate cell-to-cell fusion has been hypothesized to play roles in both pathogenesis and virus cell-to-cell propagation [51–53]. The formation of such multinucleated cells is believed to occur by the activation of SARS-CoV-2 S expressed at the cell surface of infected cells [54] and fusion with neighboring cells expressing ACE2 and serine proteases. However, the relative low abundance of ACE2+ and TMPRSS2+ cells in the lower airways suggests that there may be additional factors able to activate S [55–58]. Serine protease-independent SARS-CoV-2 S-mediated cell-cell fusion has been previously observed in vitro and reported by other studies [16, 17]. Here we show that metalloproteinases, including MMP2 and MMP9, are critical factors in SARS-CoV-2 S serine protease-independent fusion (Fig.1E,4,5). More precisely, knockdown of MMP2 and MMP9 reduced S-mediated syncytia formation in the absence of serine proteases (Fig.4). In addition, Alpha infection and replication in cells expressing high levels of MMP2 and MMP9 led to substantial fusion of infected cells that was abrogated by broad-spectrum MMP inhibitors (Fig.5). Several studies have demonstrated increased levels of MMP2 and MMP9 in patients with severe COVID-19, which associated with disease outcome [20, 22, 59, 60]. In addition, infiltrating and activated neutrophils, which are potent source of released MMP-9, are both associated with severe COVID-19 [1, 61–64]. Our studies suggest a model by which inflammation and MMP production during COVID-19 increase viral cell-to-cell transmission and exacerbate virus-induced cytopathic effects further contributing to disease.

We demonstrated that entry of SARS-CoV-2 lentiviral particles, virus-like particles and authentic Alpha variant into host cells can occur in a metalloproteinase-dependent manner (Fig.2E,3E,5B). This previously unrecognized entry pathway depended on S1/S2 processing in the viral producer cells (Fig.2,3, 6). Therefore, this third entry route is specific to SARS-CoV-2 and cannot be used by SARS-CoV-1, as SARS-CoV-1 S does not contain the critical S1/S2 furin cleavage site. Whether this additional entry pathway unique to SARS-CoV-2 played a role in its high transmissibility remain to be determined. Multiple variants of SARS-CoV-2 have emerged since the beginning of the COVID-19 pandemic, with each having their own sets of mutations that enhance immune escape and transmissibility. Interestingly, variants such as Alpha, Delta, Kappa and more recently Omicron possess mutations within the S1/S2 furin cleavage site. Notably, Delta S harbors the P681R mutation which improves S1/S2 processing and fitness over that of the ancestral virus and Alpha, which possesses the P681H mutation [12, 65–67]. Other studies, including ours, have shown that Omicron S is less efficiently processed at the S1/S2 junction compared to ancestral S, S with a D614G mutation (Fig.6B), and those of VOCs such as Delta [68, 69]. Therefore, as expected, we found that Omicron S mediated reduced metalloproteinase-dependent cell-cell fusion (Fig.6A). In addition, VLPs expressing Omicron S were only slightly sensitive to metalloproteinase-dependent entry. Omicron S has two mutations near the furin cleavage site, N679K and P681H, however the mechanism by which these or other mutations alter S processing remains to be determined. In addition, whether the inefficient use of the metalloproteinase pathway for activation of S to mediate viral entry and cell-cell fusion plays a role in the apparent distinct clinical manifestations and tropism of Omicron is unclear [11, 70–73]. Nonetheless, our findings of an additional entry pathway suggest a potential for increased tropism in the presence of MMPs during inflammation to cells that do not express serine proteases and could play important roles in dissemination and disease severity.

While this manuscript was in preparation, Yamamoto et al. reported in a pre-print a SARS-CoV-2 S-mediated metalloproteinase-dependent entry pathway in which ADAM10 was partially involved in different cell lines [74]. Although we have not directly studied a role for ADAM10 and it is still unknown if ADAM10 can cleave SARS-CoV-2 S, our findings and those of Yamamoto and colleagues highlight the promiscuity of SARS-CoV-2 for host protease activation of S and intensifies the hurdles in the usage of host protease inhibitors for therapeutic purposes. Interestingly, unlike S activation mediated via cathepsins or serine proteases, the S activation triggered via metalloproteinases required prior S processing at the S1/S2 junction. While more work is needed to determine whether MMP2, MMP9, and other metalloproteinases directly cleave S, the requirement for a processed S suggest that metalloproteinases can only cleave at the S2’ site. Further studies are required to characterize the specific roles played by the metalloproteinases and to determine the specific S cleavage site involved in the metalloproteinase pathway.

Previous studies investigating immune signatures of severe COVID-19 unveiled vascular endothelial growth factor A (VEGF-A), a disintegrin and metalloproteinases (ADAMs) and matrix metalloproteinases (MMPs) as potential markers for severe disease progression [21, 75, 76]. Our data indicate that increased secretion of MMPs during severe COVID-19 could exacerbate S-mediated cytopathic effects such as syncytia formation. Furthermore, it could also expand tropism by allowing entry in serine protease deficient cells, and potentially even in ACE2 deficient cells promoted by shed ACE2 induced by ADAM17 activity [77]. Therefore, usage of the metalloproteinase pathway by current and future circulating SARS-CoV-2 variants could have profound implications in disease severity, outcome, and potential sequelae following recovery.

## METHODS

### Cell lines, inhibitors, and antibodies

HEK293T (ATCC), HEK293T-ACE2 (kind gift of Hyeryun Choe, Scripps Research), HT1080 cells (ATCC) and Calu3 (ATCC) were cultured in Dulbecco’s Minimum Essential Medium (DMEM) supplemented with 10% fetal bovine serum (FBS, Sigma), 100 U/mL penicillin, 100 µg/mL streptomycin, and 0.3 mg/mL L-glutamine. HT1080 cells stably expressing ACE2 were generated by infection with lentiviral particles generated with psPAX2, pMDG and pLENTI_hACE2_PURO (gift from Raffaele De Francesco (Addgene plasmid # 155295)) and selection of a polyclonal HT1080-ACE2 cells using 2 µg/mL puromycin. Cells were maintained at 37°C, 5% CO2 and 100% relative humidity.

The inhibitors Camostat mesylate, GI-254023X and Batimastat were purchased from Cayman Chemical. E64d and MMP-2/MMP-9 Inhibitor II were from MilliporeSigma and TAPI-2 from Tocris.

The monoclonal antibodies SARS-CoV-1/SARS-CoV-2 Spike Protein S2 (1A9) and SARS-CoV-1/SARS-CoV-2 Nucleocapsid (6H3) were purchased from ThermoFisher Scientific. The rabbit polyclonal Anti-GAPDH antibody were purchased from Abcam. The rabbit polyclonal Anti-HIV-1 p24 antibody was purchased from MilliporeSigma. The mouse anti-N protein antibody (clone 1C7) was purchased from Bioss Antibodies, and the rabbit anti-SARS-CoV-2 spike protein (clone 007) antibody, was purchased from Sino Biological.

### SARS-CoV-2 Spike cloning and mutagenesis

The Spike gene sequence from the severe acute respiratory syndrome coronavirus 2 isolate Wuhan-Hu-1 (NC_045512.2) was codon optimized (GeneArt, ThermoFisher) and gene blocks with overlapping sequences were synthesized by Bio Basic Inc (Markham, Ontario, Canada). The full gene, untagged or with a N-terminal FLAG tag, was reconstituted by Gibson assembly, amplified by PCR, and cloned in pCAGGS. Untagged S mutants (D614G, R815A, and Δ Furin site (deletion of arginine 682, 683 and 685), Δ675-679, and variants (Alpha, Delta, Omicron) were generated by overlapping PCR and described elsewhere [78–81].

### Soluble ACE2 expression and purification

FreeStyle 293F cells (Invitrogen) were grown in FreeStyle 293F medium (Invitrogen) to a density of 1 x 10^6^ cells/mL at 37°C with 8 % CO2 with regular agitation (150 rpm). Cells were transfected with a plasmid coding for His(8)Tagged-ACE2 ectodomain (residues 1-615; [82]) using ExpiFectamine 293 transfection reagent, as directed by the manufacturer (Invitrogen). One week later, cells were pelleted and discarded. Supernatants were filtered using a 0.22 µm filter (Thermo Fisher Scientific). The soluble ACE2 (sACE2) was purified by nickel affinity columns, as directed by the manufacturer (Invitrogen). The sACE2 preparations were dialyzed against phosphate-buffered saline (PBS) and stored in aliquots at -80°C until further use. To assess purity, recombinant proteins were resolved by SDS-PAGE and stained with Coomassie Blue.

### Fusion assays

For the syncytium formation assay HEK293T and HEK293T-Ace2 cells were seeded in 24-well plates and grown to approximately 80% confluency. Cells were then transiently transfected with plasmid DNA encoding LTR-GFP (kind gift of James Cunningham, Brigham and Women’s Hospital, Boston), FLAG-SARS-CoV2 S wt or indicated mutants, and TMPRSS2 or pCAGGS in a 1:2:5 ratio using jetPRIME (Polyplus-transfection). Simultaneously, cells were placed in fresh complete media (DMEM supplemented with 10% FBS, 100 U/mL penicillin, 100 µg/mL streptomycin, 0.3 mg/mL L-glutamine) with 25 µM of Camostat or vehicle control DMSO. 24 hours post transfection, cells were imaged for syncytium formation using a ZOE Fluorescent Cell Imager (Bio-Rad) and three different fields for each well were obtained.

Cell-cell fusion assay with soluble ACE2, effector HEK293T cells were transiently transfected with plasmid DNA encoding mCherry, and SARS-CoV2 spike and target HEK293T cells were transiently transfected with plasmid DNA encoding LTR-GFP, TMPRSS2 or pCAGGS. 24 h post-transfection, effector and target cells were resuspended with 0.56 mM EDTA in PBS and co-cultured in complete media at a 1:1 ratio in the presence of increasing concentrations of soluble ACE2 (0, 25, 50, 100, 150 µg/mL). Cells were imaged 10h post-co-culture using a ZOE Fluorescent Cell Imager (Bio-Rad).

For the ZipVenus complementation cell fusion assay, HEK293T cells were seeded in a 12-well microplate (500,000 cells/well) in complete media for 24h. Transient transfections were performed using JetPRIME (Polyplus transfection, France) according to the manufacturer’s instructions. Target cells were transfected with ZipV1 (0.5µg) alone or with hACE2/pcDNA3 (0.05µg) with or without TMPRSS2/plX307(0.45µg). Effector cells population were transfected with ZipV2 (0.5µg) and SARS-CoV-2-S (0.125µg). Total DNA was normalized using the empty pCAGGS vector DNA to 1µg. Following transfection, cells were incubated at 37 °C for 24 h. Then, cells were rinsed with PBS and detached with versene (PBS, 0.53mM EDTA) and counted. 40,000 cells/well of both populations were co-seeded in complete DMEM without phenol red in a 384-well black plate with optical clear bottom and incubated for 3 hours at 37 °C, 5% CO2. Bimolecular fluorescence complementation (BiFC) signal was acquired using Biotek Synergy Neo2 plate reader (BioTek) using monochromator set to excitation/emission of 500 and 542 nm. The original BiFC constructs GCN4 leucine zipper-Venus1 (ZipV1) and GCN4 leucine zipper-Venus2 (ZipV2) were sourced from Stephen W. Michnick [83].

### RNA extraction, quantitative reverse transcription PCR, and analysis

Total RNA was extracted using RNeasy Mini Kit (Qiagen, 74104) according to manufacturer’s instructions. RNA concentrations were determined using Thermo Scientific™ NanoDrop 2000. Complementary DNA was synthesized using iScript™ Reverse Transcription Supermix (Bio-Rad, 1708840) and qRT-PCR analysis was performed using SYBR Green master mix (Life Technologies) on a Bio-Rad CFX96™ RT-PCR system. Primer sequences are shown in Supplementary Table 1.

### Gelatin zymography

HEK293T, HEK293T-ACE2, Calu3 and HT1080 cells were analyzed for MMP2 and MMP9 activity through zymographic analysis. Cells were plated in a 6-well plate, at 70-80% confluency, media was changed for FBS-free DMEM (conditioned media). Conditioned media were collected after 24h, centrifuged to remove debris and concentrated 10X using Amicon Ultra-Centrifugal filter units with a 10kD cutoff (MilliporeSigma). Total protein concentration was measured using BCA. 40µg of protein per sample were diluted in a non-reducing sample buffer (4% SDS, 20% glycerol, 0.01% bromophenol blue, 125 mM Tris-HCl, pH6.8) and loaded to a 10% Zymogram Plus (Gelatin) protein gel from Invitrogen. Following electrophoresis, gels were washed twice for 30 min in washing buffer (2.5% Triton X-100, 50mM Tris-HCl, pH7.5, 5mM CaCl_2_ and 1µM ZnCl_2_) at room temperature with gentle agitation. Gels were next incubated overnight at 37 degrees in development buffer (1%Triton X-100, 50mM Tris-HCl, pH7.5, 5mM CaCl_2_ and 1µM ZnCl_2_) to initiate enzymatic activity. Gels were stained with Coomassie Blue 0.5% for 1h and distained with 10% acetic acid and 40% methanol before being scanned.

### Lentiviral pseudotype production and entry assays

HEK293T cells were transiently co-transfected with lentiviral packaging plasmid psPAX2 (gift from Didier Trono, addgene #12260), lentiviral vector encoding LacZ or luciferase, and a plasmid encoding the viral glycoprotein (CoV S or VSV G) at a 1:1:1 ratio using jetPRIME transfection reagent. The supernatant was harvested at 48, 72, and 96 h post-transfection and filtered with a 0.45 µM filter. Lentivirus particles were concentrated via ultra-centrifugation (20,000 RPM, 1.5h, 4°C) with a sucrose cushion (20% w/v). Viral particles were resuspended with PBS and stored at -80° C.

Cells were seeded in 96-well plates to achieve approximately 50% confluence after 24 h. After 24 h, cells were pre-incubated with the inhibitor(s) for 1 h diluted in the respective standard growth media with 5 µg/mL polybrene. Concentrated lentiviruses were also diluted in growth media containing polybrene to achieve between 100-200 foci following infection. After 24 h incubation with virus and inhibitors, cells were placed in fresh growth media. 72 h post-infection, cells were fixed in formalin and stained with 100 µM X-Gal in staining solution (5 mM potassium ferrocyanide, 2 mM magnesium chloride in PBS) and incubated at 37° C for 16-24 h. Positive foci were manually counted using a light microscope. Inhibitor focus-forming units (FFUs) were normalized to vehicle control.

### Viral-like particle production and entry assays

SARS-CoV-2 virus-like particles (VLPs) were produced in HEK293T cells by co-transfection of CoV-2-N (1), CoV-2-M-IRES-E (0.5), CoV-2-Spike (0.0125) and Luc-PS9 (1) [40] at indicated ratios using jetPRIME transfection reagent (CoV-2-N, CoV-2-M-IRES-E and Luc-PS9 were gifts from Abdullah M. Syed and Jennifer A. Doudna, Gladstone Institute of Data Science and Biotechnology). N protein harboring the R203M substitution was used to enhance assembly and production of VLPs, as previously described [84]. For the bald control, the empty vector plasmid, pCAGGS, was transfected instead of the CoV-2-Spike at similar ratio. Media was changed 24 hours post-transfection and supernatants were collected at 48, 72, and 96 h post-transfection and filtered with a 0.45 µM filter. VLPs in supernatants were concentrated as described above for the lentivirus particles.

For VLP infection, cells were seeded in 96-well plates to achieve approximately 70% confluence the following day. After 24 h, cells were pre-incubated with the inhibitor(s) for 1 h diluted in 2% serum growth media with 5 µg/mL polybrene. Concentrated VLPs were also diluted in 2% serum growth media containing polybrene. After 20-24 h incubation with VLP and inhibitors, supernatant was removed, and cells were rinsed in 1X PBS and lysed by the addition of 40 µl passive lysis buffer (Promega) followed by one freeze-thaw cycle. A Synergy Neo2 Multi-Mode plate reader (BioTek) was used to measure the luciferase activity of each well after the addition of 50-100 µl of reconstituted luciferase assay buffer (Promega). Inhibitors were normalized to vehicle control.

### Knockdown of MMP2 and MMP9

293T-ACE2 and HT1080-ACE2 cells were seeded in 24-well plates and 6-well plates respectively to achieve 70% confluency after 4-6 hours and then transfected with Lipofectamine RNAiMAX (Thermofisher) using the indicated dsiRNAs (IDT, Table S1) at a final concentration of 10nM. Combination of dsiRNA was performed using a 1:1 ratio to obtain a final concentration of 10nM. After 20 hours, 293T-ACE2 were used to perform the syncytia assay as described above, and HT1080-ACE2 media was changed to conditioned media for gelatin zymography as described above. Time-course imaging of the syncytia formation was performed using an Incucyte-Zoom (EssenBioscience), and images were analyzed in imageJ to measure the percentage of green surface area over background.

### Immunoblots

Cells were washed in PBS and then lysed in cold lysis buffer (1% Triton X-100, 0.1% IGEPAL CA-630, 150mM NaCl, 50mM Tris-HCl, pH 7.5) containing protease and phosphatase inhibitors (Cell Signaling). Proteins in cell lysates were resolved by SDS-PAGE and transferred to polyvinylidenedifluoride (PVDF) membranes. Membranes were blocked for 1h at RT with blocking buffer (5% skim milk powder dissolved in 25mM Tris, pH 7.5, 150mM NaCl, and 0.1% Tween-20 [TBST]). Blots were washed in TBST and proteins were detected using the indicated primary antibodies, HRP-conjugated secondary antibodies, and visualized using chemiluminescence according to manufacturer protocol (Bio-Rad Clarity ECL substrate).

### Microneutralization assay using live SARS-CoV-2 alpha (B.1.1.7) variant

A previously described in vitro microneutralization assay [85, 86] was performed with modifications and using the SARS-CoV-2 alpha variant (B.1.1.7 lineage). HT1080-ACE2 cells were cultured in DMEM supplemented with penicillin (100 U/mL), streptomycin (100 µg/mL), HEPES, L-Glutamine (0.3 mg/mL), 10% FBS (all from Thermo Fisher Scientific) and puromycin (1 μg/mL, InvivoGen). Twenty-four hours before infection, 2.5×10^4^ HT1080 ACE2 cells were seeded per well of duplicate 96 well plates in puromycin-deficient DMEM and cultured overnight (37°C/5% CO_2_) for cell monolayer to adhere. On the day of infection, a deep well plate was used to perform 1:2 serial dilutions for inhibitors listed below in MEM supplemented with penicillin (100 U/mL), streptomycin (100 µg/mL), HEPES, L-Glutamine (0.3 mg/mL), 0.12% sodium bicarbonate, 2% FBS (all from Thermo Fisher Scientific) and 0.24% BSA (EMD Millipore Corporation). The inhibitors Camostat (range: 40 μM – 1.25 μM), TAPI-2 (range: 20 μM – 0.675 μM), GI 254023X (range: 40 μM – 1.25 μM) and E64d (range: 20 μM – 1.25 μM) were included in this assay. MEM + 2% FBS containing DMSO at an equivalent concentration to the above inhibitor dilutions served as the vehicle control. All media was aspirated from 96 well plates seeded with HT1080-ACE2 cells and replaced with 100 μL appropriate inhibitor dilution (or vehicle control). Promptly, 2×10^3^ TCID50/mL SARS-CoV-2 alpha variant was prepared in a Biosafety Level 3 laboratory (ImPaKT Facility, Western University) and a volume corresponding to 100 TCID50 virus per well was added to wells already containing inhibitor or vehicle diluted in media. An equivalent volume of media void of virus was added to uninfected control wells. All wells were gently mixed and cultured overnight at 37°C/5% CO_2_.

After overnight culture, media was discarded and replaced with 10% formaldehyde for >24 hours to cross-link cell monolayers. Wells were washed with PBS, permeabilized for 15 minutes with PBS + 0.1% Triton X-100 (BDH Laboratory Reagents), washed again in PBS and then blocked for one hour with PBS + 3% non-fat milk. At this point, one plate was processed as detailed previously [85] to quantify virus infection. Briefly, a mouse anti-SARS-CoV-2 nucleocapsid (N) protein primary antibody and an anti-mouse IgG HRP secondary antibody in conjunction with SIGMAFAST™ OPD developing solution (Millipore Sigma) permitted SARS-CoV-2 infection quantification. The optical density at 490 nm served as the assay readout and was measured using a Synergy LX multi-mode reader and Gen5 microplate reader and imager software (Agilent).

In parallel and after blocking, the second plate was incubated for one hour with a primary antibody solution formulated in PBS + 1% non-fat milk containing both mouse anti-N protein (1 μg/mL, clone 1C7) and rabbit anti-SARS-CoV-2 spike protein (1:500 dilution, clone 007) antibodies. Extensive washing with PBS ensued, followed by a 45-minute incubation with donkey anti-mouse IgG Alexa Fluor Plus 488 (1 μg/mL, Invitrogen), donkey anti-rabbit IgG Alexa Fluor Plus 594 (2 μg/mL, Invitrogen) antibodies and DAPI (1:1000, Millipore Sigma) in PBS + 0.5% BSA consisting of. All wells were then washed three times in PBS, monolayers were covered with minimal PBS and fluorescence images were acquired with an EVOS™ M7000 Imaging System (Invitrogen).

### Statistical analysis

Data are expressed as mean ± standard deviation of the mean (SD). Significance was determined by analysis of variance (one-way ANOVA) followed by a Dunnett’s multiple comparisons test. A p-value lower than 0.05 was used to indicate a statistically significant difference ****, *P* <0.0001, ***, *P* <0.001,**, *P* <0.01, *P* *<0.05. Statistical analyses were performed with GraphPad Prism 9.

## ACKNOWLEDGEMENTS

We would like to acknowledge technical support from the uOttawa Flow Cytometry & Virometry Core Facility and the uOttawa Cell Biology and Image Acquisition Core Facility. We would also like to acknowledge support from the Western University ImPaKT staff for maintaining the facility needed for our work.

This research was funded by a Bhagirath Singh Early Career Award in Infection and Immunity to M.C., a COVID-19 Rapid Research grant from the Canadian Institutes for Health Research (CIHR, OV3 170632), and CIHR stream 1 for SARS-CoV-2 Variant Research to M.C. and P.G. Part of this research was also supported by CIHR operating Pandemic and Health Emergencies Research grant #177958, a CIHR stream 1 and 2 for SARS-CoV-2 Variant Research to A.F. Part of this research was also supported by CIHR Operating Grant: Emerging COVID-19 Research Gaps and Priorities #466984 to S.M. This work was also supported by the Sentinelle COVID Quebec network led by the Laboratoire de Santé Publique du Québec (LSPQ) in collaboration with Fonds de Recherche du Québec-Santé (FRQS) and Genome Canada – Génome Québec, and by the Ministère de la Santé et des Services Sociaux (MSSS) and the Ministère de l’Économie et Innovation (MEI). Funding was also provided by an operating grant from CIHR from the Canadian 2019 Novel Coronavirus (COVID-19) Rapid Research Funding Opportunity (FRN440388 to JDD and GAD) and an Infrastructure Grant from CFI for the Imaging Pathogens for Knowledge Translation (ImPaKT) Facility (#36287 to JDD and GAD).

K.F. and R.P.M. were supported by Ontario Graduate Scholarships (OGS). C.M.S. was supported by a graduate scholarship from the Natural Sciences and Engineering Research Council of Canada and an OGS. M.C. is a Canada Research Chair in Molecular Virology and Antiviral Therapeutics (950-232840) and a recipient of an Ontario Ministry of Research, Innovation and Science Early Researcher Award (ER18-14-09). A.F. is a Canada Research Chair on Retroviral Entry (RCHS0235 950-232424). The funders had no role in study design, data collection and analysis, decision to publish, or preparation of the manuscript.

## AUTHORS CONTRIBUTIONS

GAD, JDD and MC conceived the study. MB, GL, CF, KF, RPM, AP, AA, CMS, GAD, JDD, and MC designed experimental approaches. MB, GL, CF, KF, RPM, AP, AA, CMS, JP, GBB, RD, YB, JYL, MC performed experiments. WLS, SM, AF provided resources. MB, GL, CF, KF, RPM, AP, AA, GAD, JDD, MC analyzed and interpreted results. PMG, GAD, JDD, MC supervised the study. MB, GL, MC wrote the original draft. Every author has read and edited the manuscript

## EXPANDED VIEW FIGURE LEGENDS

**Supplemental Figure 1.**
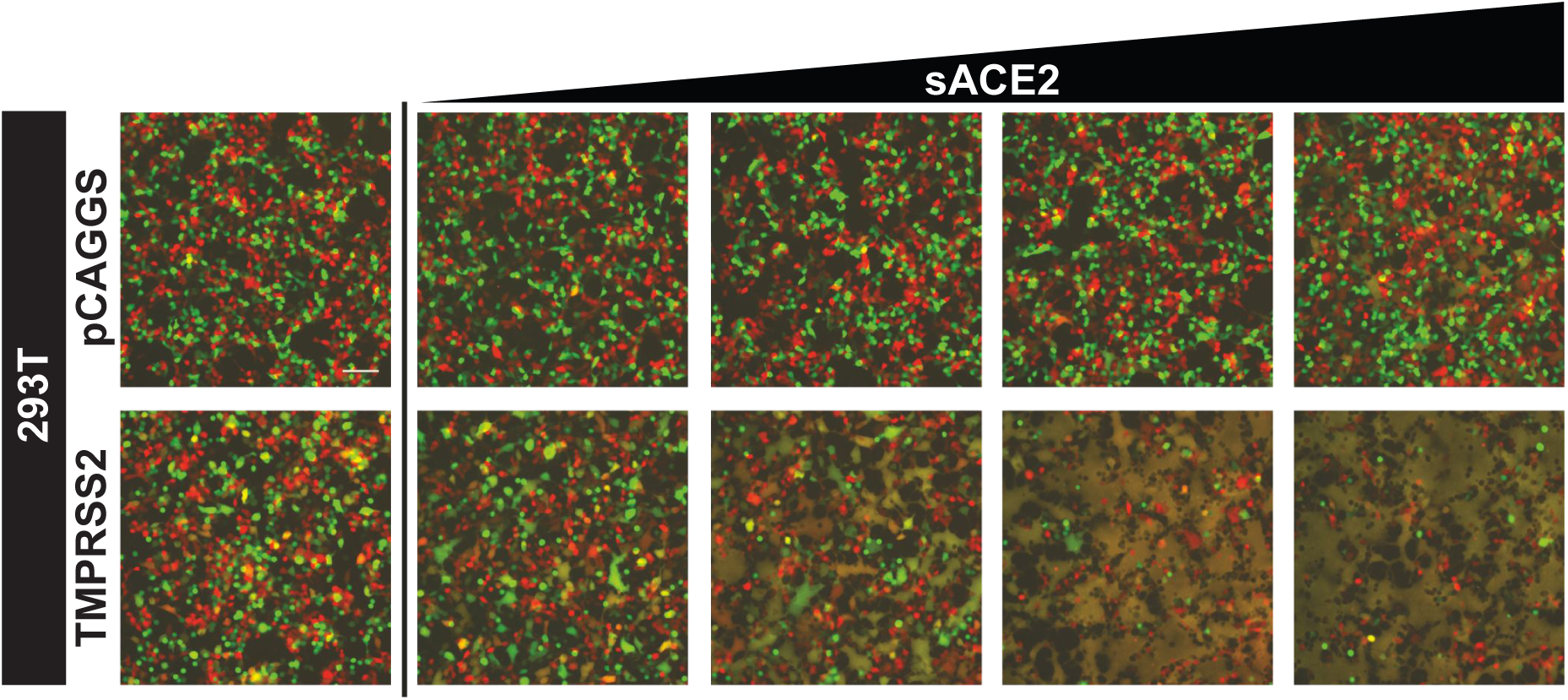
TMPRSS2 enhances SARS-CoV-2 S fusion activity in an ACE2 dose-dependent manner. HEK293T cells expressing mCherry and SARS-CoV2 spike were co-cultured with HEK293T cells expressing GFP and TMPRSS2 or with HEK293T cells expressing GFP only in the presence of increasing concentrations of soluble ACE2 (0, 25, 50, 100, 150 µg/mL). Cells were imaged 10h post-co-culture. Representative images are shown.

**Supplemental Figure 2.**
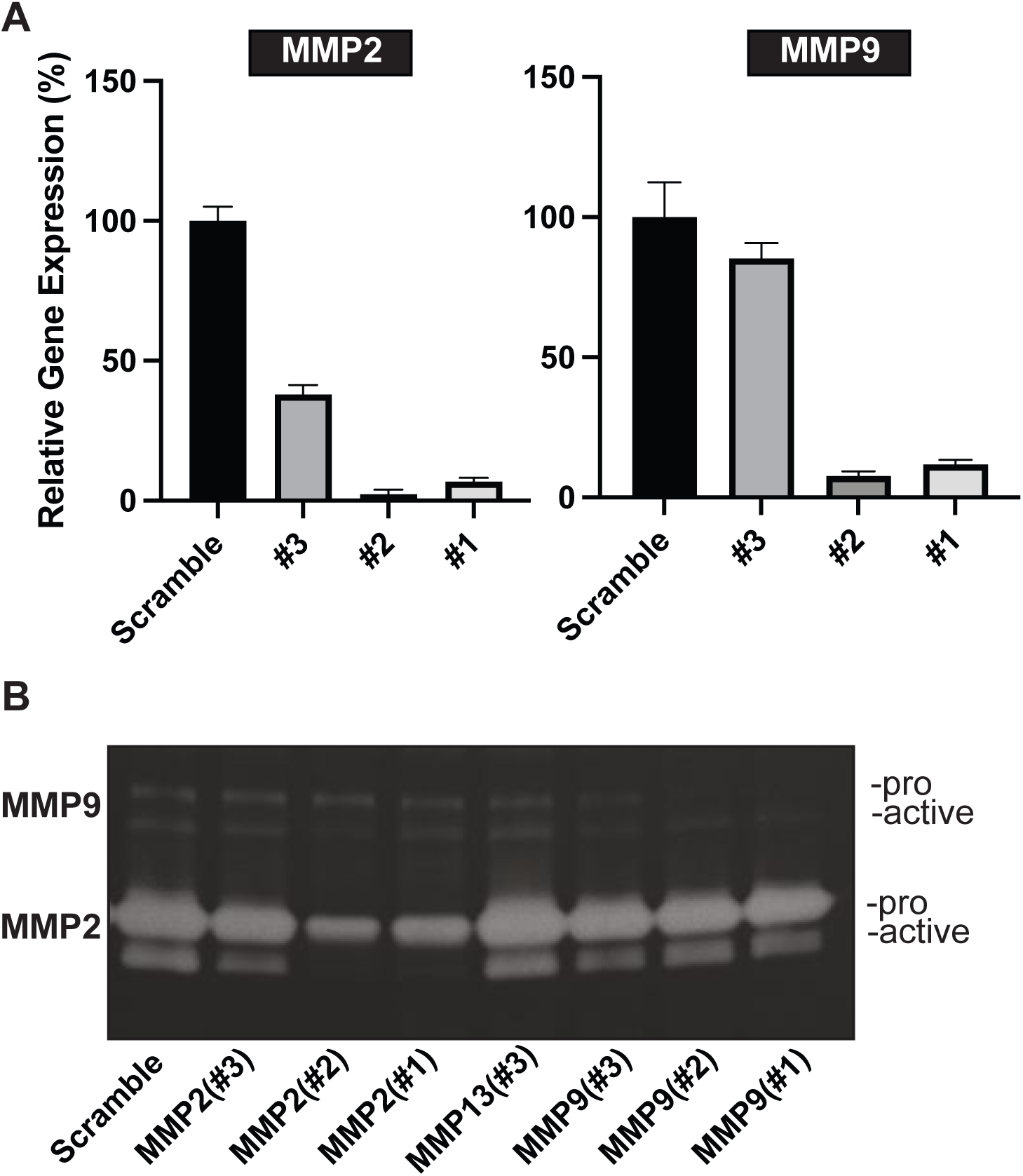
MMP2/MMP9 knockdown reduces production and secretion in HT1080-ACE2 cells. **(A)** HT1080-ACE2 cells were transfected with the indicated dsiRNAs at a final concentration of 10nM, and relative mRNA levels of MMP2 and MMP9 was measured by RT-qPCR. The level of actin mRNA expression in each sample was used to standardize the data, and normalization on scramble genes expression was performed. **(B)** Gelatin zymogram of conditioned media (24h) from HT1080-ACE2 cells in **(A),** arrows indicate the pro- and active- MMP2 or MMP9.

**Supplemental Figure 3.**
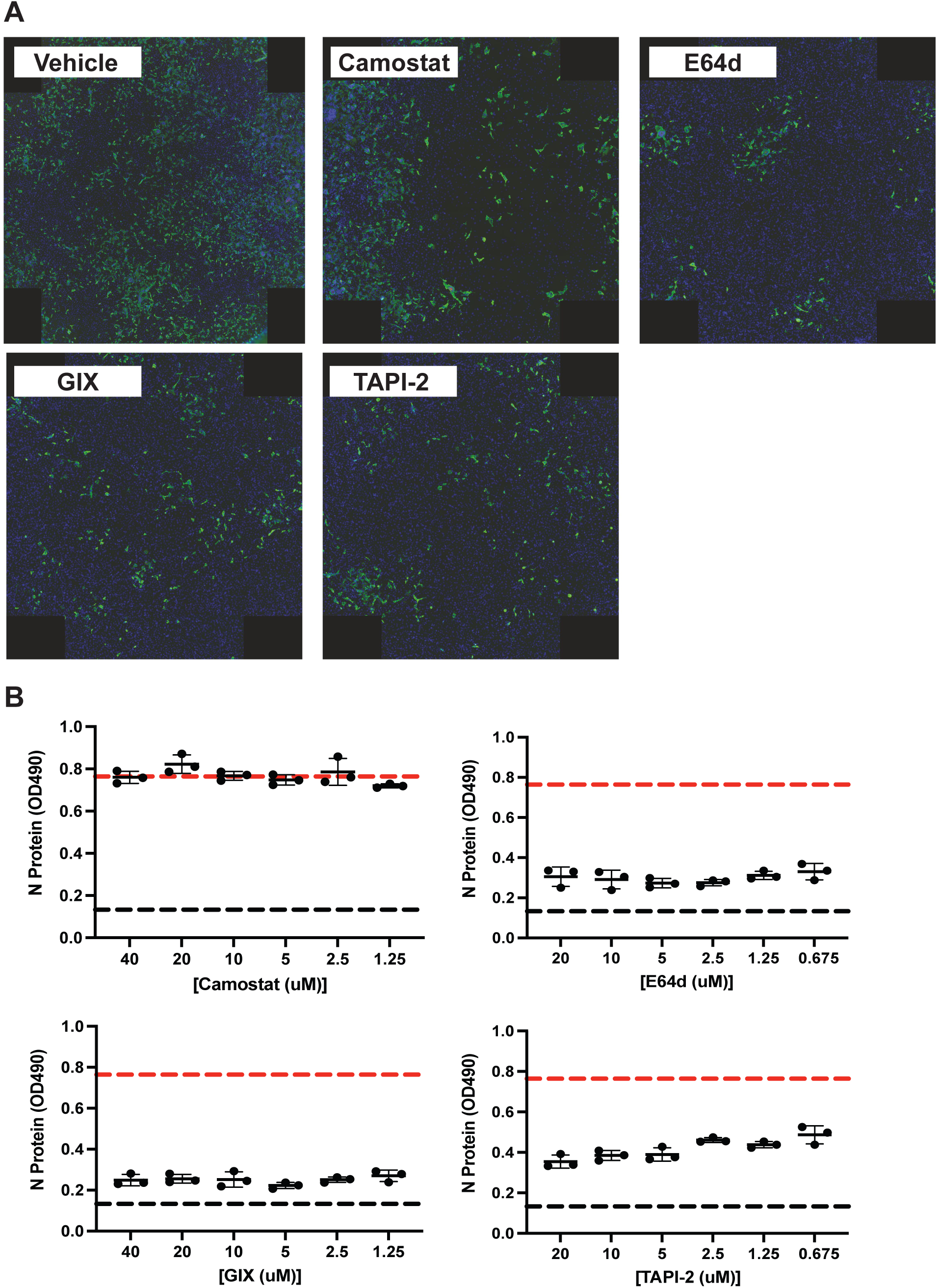
Syncytia formation and infection by replicative Alpha are blocked by metalloproteinase inhibitors in HT1080-Ace2 cells. **(A**) Visualization of HT1080-ACE2 syncytia formation after infection by Alpha in presence of indicated inhibitors or vehicle. Cells were treated with Camostat (20 uM), E64D (10 uM), TAPI-2 (20 uM), GIX (10 uM) or Vehicle (DMSO), following prompt addition of Alpha variant. After 20h, cells were washed, blocked, and stained with rabbit anti-SARS-CoV-2 spike (S), mouse anti-SARS-CoV-2 nucleocapsid (N) followed by staining with DAPI, donkey anti-mouse IgG Alexa Fluor Plus 488 and donkey anti-rabbit IgG Alexa Fluor Plus 594 antibodies. Nuclei, S and N proteins are shown in purple, red and green respectively. Fluorescent images were acquired with an EVOS™ M7000 Imaging System. Images are representative of 3 independent experiments. **(B)** SARS-CoV-2 infection quantification following infection in presence of indicated inhibitors (0.675-40uM) or vehicle. 20h post-infection, cells were washed, blocked, permeabilized and stained with mouse anti-SARS-CoV-2 N protein followed by an anti-mouse IgG HRP in conjunction with SIGMAFAST™ OPD developing solution. Optical density (OD) at 490 nm was measured using Synergy LX multi-mode reader and Gen5 microplate reader and imager software. The red line indicates the OD obtained for vehicle. Each bar shows the mean of triplicate values of 3 independent experiments with error bars showing standard deviation. Significance was determined by analysis of variance (one-way ANOVA) followed by a Dunnett’s multiple comparisons test. P-value lower than 0.05 was used to indicate a statistically significant difference (****, *P* <0.0001, ***, *P* <0.001, **, *P* <0.01, *, *P*<0.05).

**Supplemental Figure 4.**
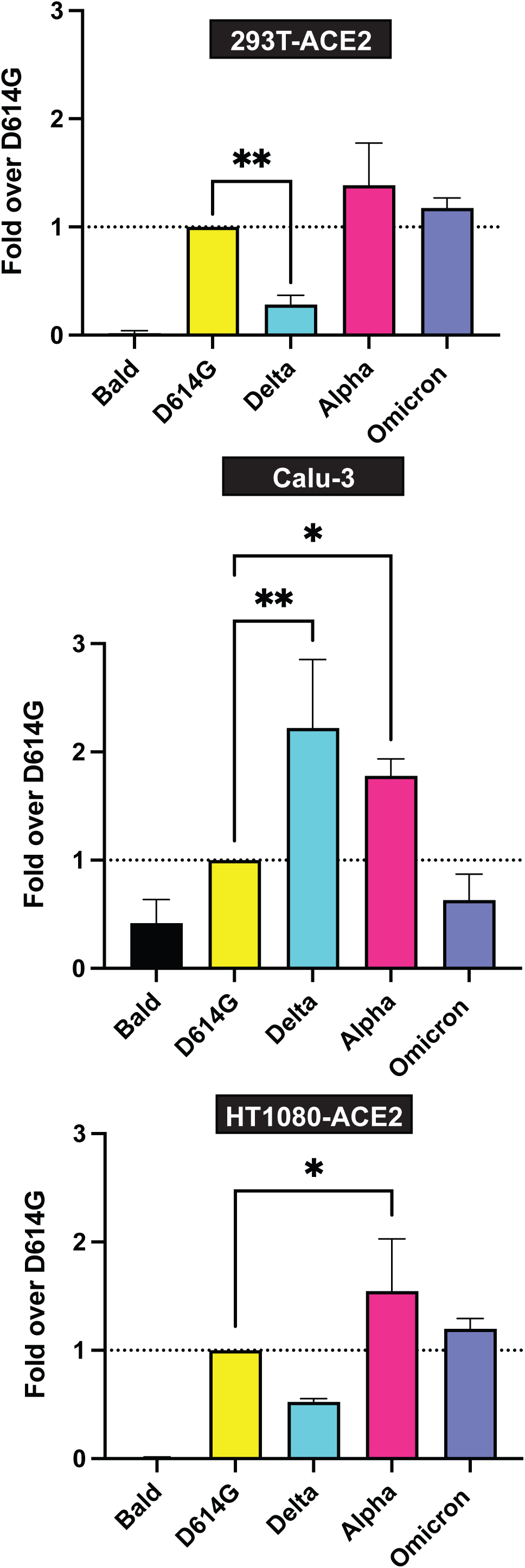
Omicron uses less efficiently the TMPRSS2-dependent entry pathway compared to other variant of concerns. Variant of concerns virus-like particle entry assay on 293T-ACE2 **(A)**, Calu-3 **(B)** and HT1080-ACE2 **(C)**. VLPs entry were measured 24h after incubation with VLP and normalized as fold over D614G. Each bar shows the mean of triplicate values of 3 independent experiments with error bars showing standard deviation. Significance was determined by analysis of variance (one-way ANOVA) followed by a Dunnett’s multiple comparisons test. P-value lower than 0.05 was used to indicate a statistically significant difference (****, *P* <0.0001, ***, *P* <0.001, **, *P* <0.01, *, *P* *<0.05)

