## Supplementary figures and images for "Identification of a SARS-CoV-2 host metalloproteinase-dependent entry pathway differentially used by SARS-CoV-2 and variants of concern Alpha, Delta, and Omicron"

### Supplemental tables

Supplementary Table 1, RT-qPCR primer sequences

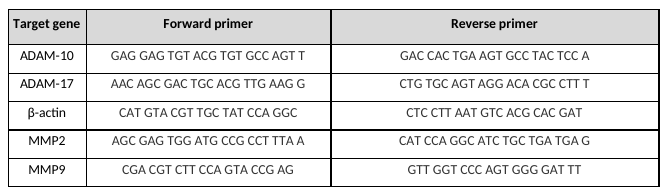

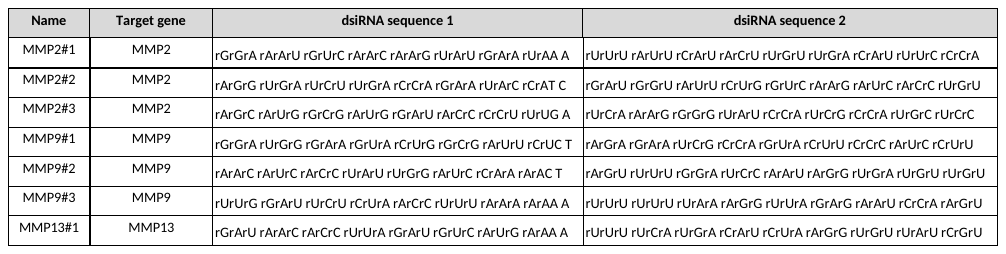
Supplementary Table 2, dsiRNA sequences
